# Blind individuals’ enhanced ability to sense their own heartbeat is related to the thickness of their occipital cortex

**DOI:** 10.1101/2024.02.26.581908

**Authors:** Anna-Lena Stroh, Dominika Radziun, Maksymilian Korczyk, Laura Crucianelli, H. Henrik Ehrsson, Marcin Szwed

## Abstract

Blindness is associated with heightened sensory abilities, such as improved hearing and tactile acuity. Moreover, recent evidence suggests that blind individuals are better than sighted individuals at perceiving their own heartbeat, suggesting enhanced interoceptive accuracy. Structural changes in the occipital cortex have been hypothesized as the basis of these behavioral enhancements. Indeed, several studies have shown that congenitally blind individuals have increased cortical thickness within occipital areas compared to sighted individuals, but how these structural differences relate to behavioral enhancements is unclear. This study investigated the relationship between cardiac interoceptive accuracy and cortical thickness in 23 congenitally blind individuals and 23 matched sighted controls. Our results show a significant positive correlation between performance in a heartbeat counting task and cortical thickness only in the blind group, indicating a connection between structural changes in occipital areas and blind individuals’ better ability to perceive heartbeats.

---

Numerous studies with blind individuals have provided evidence of the brain’s ability to adapt and change, as well as compensatory behavioral adjustments in relation to a lack of vision. For example, it has been shown that blind individuals have finer tactile discrimination thresholds (Alary et al., 2009; Goldreich & Kanics, 2003, 2006; Radziun et al., 2023a); they perform better in spatial sound localization (Collignon & De Volder, 2009; Gougoux et al., 2005; Lessard et al., 1998; Röder et al., 1999; Voss et al., 2004), auditory pitch discrimination (Gougoux et al., 2004); they have better verbal memory (Amedi et al., 2003; Pasqualotto et al., 2013; Raz et al., 2007; Röder et al., 2001), and they also seem to be better at processing, learning, and memorizing voices (Bull et al., 1983; Föcker et al., 2012). Behavioral enhancements in blind individuals could be linked to intramodal plasticity, i.e., plasticity within non-deprived auditory and somatosensory areas (see Fiehler & Rösler, 2010), and/or cross-modal plasticity, i.e., reorganization of brain areas that are typically associated with vision in sighted individuals (Amedi et al., 2003, 2004; Gougoux et al., 2005; Merabet & Pascual-Leone, 2010; Voss & Zatorre, 2012). Compelling evidence for this last notion comes from studies showing that activation of the occipital cortices is related to behavioral improvements in blind individuals. For example, several functional magnetic resonance imaging (fMRI) studies have shown that the level of occipital cortex activation is related to superior behavioral performance in verbal memory and sound localization (Amedi et al., 2003; Gougoux et al., 2005). Moreover, disruption of the occipital cortex in blind participants by means of transcranial magnetic stimulation (TMS) has been shown to impair performance in verbal memory (Amedi et al., 2004) and auditory spatial localization tasks (Collignon et al., 2009).

In addition to these functional changes, early blindness has been shown to lead to structural changes. It is now well-established that blind individuals have thicker occipital cortices than sighted individuals (Aguirre et al., 2016; Anurova et al., 2015; Bridge et al., 2009; Hasson et al., 2016; Jiang et al., 2009; Park et al., 2009; Voss & Zatorre, 2012). Jiang et al. (2009) proposed that increased cortical thickness in blind individuals was likely due to the disruption of synaptic pruning, while Park et al. (2009) suggested that it was the result of cross-modal plasticity. These two proposals are not mutually exclusive and, in fact, it could be hypothesized that the cross-modal engagement observed in blind individuals is possible because certain cortico-cortical connections are not pruned but rather preserved and strengthened (see Singh et al., 2018). Nevertheless, relating these structural changes to specific behavioral enhancements has proven rather elusive. So far, only Voss and Zatorre (2012) have managed to relate the cortical thickness of occipital areas to behavioral enhancements in blind individuals in pitch discrimination and musical tasks. However, the correlations between cortical thickness and behavioral measures they reported were only investigated in a blind group, without a sighted control group. The evidence would have been stronger if it had been possible to demonstrate that these correlations were specific to the blind group and were not present in the sighted group.

So far, most studies have focused on behavioral enhancements that facilitate blind people’s interaction with the external environment. Recently, however, Radziun et al. (2023b) have shown that such enhancements may extend to interoception, i.e., a group of sensations arising from one’s internal organs that convey information about the physiological state of the body (Khalsa et al., 2018). Specifically, it was shown that a group of blind individuals had significantly higher accuracy in perceiving their own heartbeat than a group of sighted controls. Given the importance of interoception in a variety of vital functions, including emotional processing (Critchley & Garfinkel, 2017) and bodily self-awareness (Crucianelli et al., 2018; Herbert & Pollatos, 2012; Quigley et al., 2021), this finding opens up important research avenues in relation to the impact and extent of compensatory brain plasticity. However, the neuroanatomical basis of this interoceptive enhancement in blind individuals is unknown.

As blind individuals rely heavily on hearing and touch to interact with their environment, it seems reasonable that neurophysiological changes could manifest within brain regions responsible for auditory and somatosensory processing. This type of plasticity is also known as intramodal plasticity (e.g., De Borst & De Gelder, 2019). However, most studies investigating structural plasticity in blind individuals have not found any structural differences between blind and sighted individuals in the auditory (Noppeney et al., 2005; Pan et al., 2007; Ptito et al., 2008) or the somatosensory cortex (Li et al., 2017). While the evidence for intramodal plasticity is scarce, numerous studies have shown that blind individuals have thicker occipital cortices (Bridge & Watkins, 2019; Jiang et al., 2009; Park et al., 2009) than sighted individuals. Crucially, blind individuals’ behavioral enhancements in the auditory domain seem to be related to their thicker occipital cortices (Voss & Zatorre, 2012). Based on this evidence, it could be hypothesized that enhanced cardiac interoceptive accuracy in blind individuals is supported by regions of the brain that are typically associated with visual processing in sighted individuals, i.e., that cross-modal plasticity supports blind individuals’ enhanced ability to sense heartbeats.

Here, we wanted to assess whether enhanced cardiac interoception in blind individuals is related to changes in brain structure. To this end, we used structural magnetic resonance imaging (sMRI) to measure cortical thickness in a group of congenitally blind individuals and a sighted control group, and we correlated this anatomical measure to cardiac interoceptive accuracy in the same individuals. We analyzed cortical thickness, which is an established neuroanatomical measure of great interest in both normal development and developmental plasticity and is one of the most consistently reported structural changes observed in blind individuals (Aguirre et al., 2016; Anurova et al., 2015; Bridge et al., 2009; Hasson et al., 2016; Jiang et al., 2009; Park et al., 2009; Voss & Zatorre, 2012).

## Methods

### Participants

23 blind and 23 sighted individuals matched for age, sex, and reported handedness were included in the study (age range = 22–45 years; mean age = 33.30 years; 14 men and 9 women per group). Behavioral data were collected from 22 blind participants and 12 sighted participants as part of a study by Radziun et al. (2023b). The heartbeat counting task was administered to one additional blind participant and 11 additional sighted participants. The MRI data of the blind participants were collected as a part of another project (Korczyk et al., in preparation). The MRI data of the sighted participants were collected specifically for the present study. Neuroimaging and behavioral data were available for two additional blind participants, but they were excluded from the analyses due to their failure to successfully complete the behavioral task, as described in Radziun et al. (2023b). The demographic data of the final sample are summarized in Table 1.

**Table 1.** Blind participant characteristics.

| Participant | Age (years) | Sex | Cause of blindness | Reading hand (finger) | Age when learned Braille | Reading frequency |
| --- | --- | --- | --- | --- | --- | --- |
| 1 | 32 | female | retinopathy of prematurity | right (index finger) | 7 | rarely |
| 2 | 45 | female | retinopathy of prematurity | right (index finger) | 7 | every day |
| 3 | 42 | male | toxoplasmosis | right (index finger) | 8 | often |
| 4 | 32 | female | retinopathy of prematurity | left | 6 | rarely |
| 5 | 31 | male | retinopathy of prematurity | right | 5 | once a week |
| 6 | 40 | female | retinopathy of prematurity | left | 6 | none |
| 7 | 35 | male | glaucoma | right | 7 | often |
| 8 | 24 | male | atrophy of the optic nerve | left (index finger) | 6 | every day |
| 9 | 22 | male | microphthalmia | left | 4 | every day |
| 10 | 30 | female | atrophy of the optic nerve | right | 4 | rarely |
| 11 | 30 | male | optic nerve hypoplasia | right | 7 | once a week |
| 12 | 35 | female | undefined (genetic) | right (index finger) | 4 | rarely |
| 13 | 34 | male | undefined (genetic) | left | 7 | every day |
| 14 | 27 | male | retinopathy of prematurity | right | 7 | rarely |
| 15 | 28 | female | retinopathy of prematurity | right | 8 | every day |
| 16 | 31 | female | atrophy of the optic nerve | right (index finger) | 7 | often |
| 17 | 25 | male | retinopathy of prematurity | left | 7 | rarely |
| 18 | 45 | male | retinopathy of prematurity | left | 7 | rarely |
| 19 | 26 | male | retinopathy of prematurity | left | 7 | every day |
| 20 | 43 | male | atrophy of the optic nerve | left (index finger) | 7 | rarely |
| 21 | 39 | male | retinopathy of prematurity | left | 5 | rarely |
| 22 | 39 | female | retinopathy of prematurity | left (index finger) | 6 | often |
| 23 | 31 | male | retinopathy of prematurity | left | 6 | once a week |

For all blind participants, blindness was attributed to a peripheral origin and was not associated with any other sensory disabilities. To be included in the study, participants had to either be completely blind or, at most, have minimal light sensitivity that did not allow them to use this sense in a functional way, and they could not have any pattern vision. All sighted participants had normal or corrected-to-normal vision. None of the participants reported any history of psychiatric or neurological conditions.

All participants gave written informed consent and received monetary compensation for their participation. The experiment was approved by the local ethics committee of Jagiellonian University.

### Experimental tasks and procedure

#### Behavioral tasks

According to Garfinkel and colleagues’ (2015) dimensional model of interoception, three major dimensions of interoception can be distinguished: (1) interoceptive accuracy, meaning behavioral performance on a test consisting of monitoring one’s own physiological events (here, the heartbeat counting task; Schandry, 1981); (2) interoceptive sensibility, meaning the participant’s assessment of their own interoceptive experiences, as obtained by self-report (here, Multidimensional Assessment of Interoceptive Awareness questionnaire; Mehling et al., 2012); (3) interoceptive awareness, meaning the degree to which interoceptive accuracy correlates with confidence in task response (the relationship between the accuracy in the heartbeat counting task and the level of confidence reported by the participant after each trial). Crucially, since interoceptive accuracy was found to be significantly higher in blind participants compared to sighted volunteers in our previous behavioral study (Radziun et al., 2023b) and is the measure that most directly reflects the ability to sense heartbeats, we focused our structural MRI analyses on interoceptive accuracy.

First, the participants were asked to fill out the Multidimensional Assessment of Interoceptive Awareness questionnaire (MAIA; Mehling et al., 2012; see Brytek-Matera & Kozieł, 2015 for a Polish translation and validation). The MAIA consists of 32 items that cover eight distinct dimensions of body perception: Noticing, Not-Distracting, Not-Worrying, Attention Regulation, Emotional Awareness, Self-Regulation, Body Listening, and Trusting. The questionnaire has a range of scores of 0–5, with 0 indicating low and 5 indicating high interoceptive sensibility. As previous research has shown that heightened physiological arousal can enhance the perception of heartbeats (see Pollatos et al., 2007), the questionnaire was administered to participants at the start of the study rather than at the end. We implemented this process to ensure that any possible increase in heart rate due to factors such as walking briskly to the study location could return to a normal level. For the same reason, the participants were instructed not to consume any beverages that contained caffeine on the day of the study (Hartley et al., 2004; McMullen et al., 2012).

Before starting the heartbeat counting task, the participants were provided with an overview of the experimental procedure and given a brief outline of what would occur next. Each participant sat on a chair in a comfortable position. Prior to the heartbeat counting task, a baseline heart-rate recording was taken over a 5-minute period. We measured the participants’ heart rate using a Biopac MP150 BN-PPGED (Goleta, CA, United States) pulse oximeter, attached to the left index finger. The device was connected to a laptop running AcqKnowledge software (version 5.0), which recorded the number of heartbeats. To prevent participants from sensing their pulse in their fingers due to the pulse oximeter’s grip, we carefully adjusted the finger cuff to be comfortably snug without being too tight (see Murphy et al., 2019).

The number of heartbeats was quantified using the embedded ‘count peaks’ function of the AcqKnowledge software. Sighted volunteers were blindfolded while performing the tasks (see Radziun et al., 2023b). The participants were given the following instructions: “Without manually checking, can you silently count each heartbeat you feel in your body from the time you hear ‘start’ to when you hear ‘stop’? Do not take your pulse or feel your chest with your hand. You are only allowed to feel the sensation of your heart beating” (adapted from Garfinkel et al., 2015). After each trial, participants verbally reported the number of heartbeats counted. They did not receive any feedback regarding their performance. Immediately after providing the number of heartbeats counted, the participants were requested to assess how confident they were in the accuracy of their answers (Garfinkel et al., 2015). This confidence judgment was made using a scale ranging from 0 (total guess/no heartbeat awareness) to 10 (complete confidence/full perception of the heartbeat). A 30- second break was provided before the start of the next trial. Each participant completed six trials with a duration of 25, 30, 35, 40, 45, and 50 seconds, presented in a randomized order. No information was provided to the participants regarding the duration of the intervals.

#### MRI acquisition and image processing

MRI data were collected at Małopolskie Centrum Biotechnologii in Kraków, Poland using a 3T Siemens Skyra scanner equipped with a padded 64-channel head coil. T1-weighted images were acquired using a magnetization-prepared rapid gradient-echo (MPRAGE) sequence (TR = 1800 ms, TE = 2.37 ms, flip angle = 8⁰, field of view = 250 mm, 208 coronal slices, voxel size = 0.729 mm^3^).

The anatomical images and surface-based morphometry were processed using the recon-all function of FreeSurfer (version 7.2.0, http://surfer.nmr.mgh.harvard.edu/) with default parameter settings. A full description of the processing steps can be found elsewhere (Fischl & Dale, 2000). Briefly, image reconstruction involved intensity normalization, transformation to Talairach atlas space, and removal of non-brain tissue. The boundary between white matter and gray matter was determined using intensity, neighborhood, and smoothness constraints. Then, the space between the pial surface and white matter boundary was tessellated and smoothed to create a cortical ribbon. The cortical ribbon was parcellated, and neuroanatomical labels of brain areas were assigned to each voxel based on probabilistic anatomic information and landmarks (Dale et al., 1999; Fischl et al., 2002; Fischl et al., 2004). Cortical thickness was calculated as the closest distance between the gray/white matter boundary and the gray/pial boundary at each vertex of both hemispheres (Fischl & Dale, 2000). The left and right hemispheres of all participants were registered to the fsaverage atlas (common surface space) templates included in FreeSurfer and smoothed with a Gaussian kernel of FWHM 10 mm. Each hemisphere was modeled separately.

### Data analysis

#### Behavioral analysis

The data were tested for normality using the Shapiro–Wilk test, and the interoceptive accuracy was found to be not distributed normally (p < .05). Therefore, nonparametric statistics were used (Mann–Whitney U test for independent group comparisons). All p-values are two-tailed.

#### Interoceptive accuracy

For each participant, an accuracy score was derived, resulting in the following formula for interoceptive accuracy across all trials (Schandry, 1981):

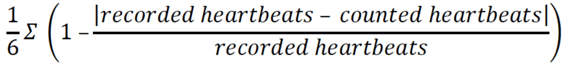

The interoceptive accuracy scores obtained using this formula usually vary between 0 and 1, with higher scores indicating better counting of the heartbeats (i.e., smaller differences between estimated and actual heartbeats). However, in instances of extreme values reported as counted heartbeats, the formula permits scores to extend from negative infinity to 1. Two blind participants were excluded from the analyses due to their failure to successfully complete the task (extremely low accuracy levels of −0.128 and −1.178; see *Participants*).

#### Interoceptive sensibility

The mean MAIA scores were used as a measure of overall interoceptive sensibility, with higher scores indicating higher interoceptive sensibility.

The average confidence level in counting heartbeats, which is another measure of interoceptive sensibility, was computed for every participant by averaging the confidence ratings across all experimental trials, resulting in an overall measure of the mean confidence in the perceived accuracy of responses.

#### Structural analysis

First, we wanted to replicate previous studies and compare cortical thickness between the groups of congenitally blind and sighted individuals by fitting general linear models at each vertex using FreeSurfer for both the left and right hemispheres. In the next step, statistical analysis was performed at each vertex to test the significance of the correlation between interoceptive accuracy and cortical thickness by including interoceptive accuracy in a separate model.

Analyses were performed over the whole brain. Given that we had a priori hypotheses about thicker visual cortical thickness in blind compared to sighted individuals and to avoid Type II errors, we additionally performed analyses that were restricted to the visual cortex and cortical areas that have been shown to be involved in cardiac interoception (Schulz et al., 2016; outlined in yellow in Figure S1 in Supplementary Material). The reconstructed cortical surface was automatically parcellated for each participant into the 180 cortical areas defined in the HCP-MMP1.0 atlas (Glasser et al. 2016). Then, early visual areas (V1, V2, V3, V4), insular areas (MI, PoI2, AAIC), cingulate area 24dd, area 43, as well as parietal area PFcm were combined to generate a mask of the visual cortex and regions that have been shown to be involved in cardiac interoception (see Schulz, 2016; see Figure 1). Cluster-wise correction for multiple comparisons was performed by running permutation tests with the mri_glmfit-sim tool provided by FreeSurfer (1000 iterations per hemisphere). The vertex-wise threshold was set to p < 0.01 (two-sided; Greve & Fischl, 2018). Statistical maps are displayed on the inflated surface of the FreeSurfer standard brain, thresholded at a vertex-wise threshold of p < 0.01.

**Figure 1.**
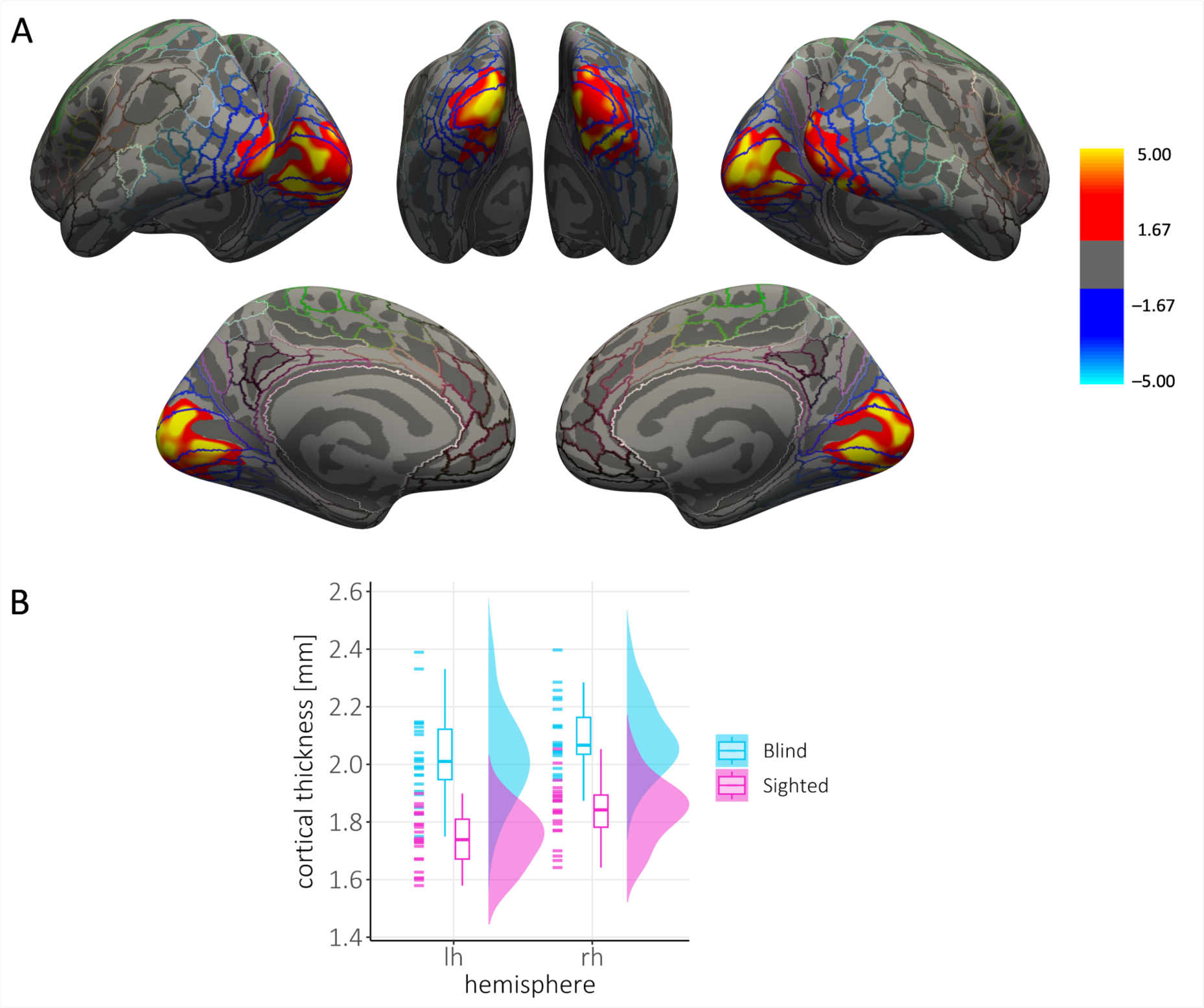
Group differences in visual cortical thickness. **A** Thresholded statistical significance maps (vertex-wise p < .01, cluster-wise p < .05, two-sided) display cortical thickness differences between congenitally blind individuals (CB group, n = 23) and sighted controls (SC group, n = 23). Maps are superimposed on the inflated surface (dark gray: sulci, light gray: gyri) of the FreeSurfer standard brain. Colored lines indicate parcellations of the HCP-MMP1.0 atlas (Glasser et al. 2016). Clusters with higher cortical thickness in the CB group are marked in red/yellow. **B** Average individual (single data points) and group mean cortical thickness extracted from significant clusters.

## Results

### Behavioral results

#### Interoceptive accuracy

Blind participants had better interoceptive accuracy than sighted participants, as reflected by significantly higher performance in the heartbeat counting task (W = 170, p = .038, CI95% = 0.007–0.227, M_Blind_ = 0.739, M_Sighted_ = 0.622). The heart rate was 75.11 BPM in the blind group and 76.93 BPM in the sighted group; there was no significant difference between the groups (W = 251.5, p = .814, CI95% = −4.000–6.000).

#### Interoceptive sensibility

There was no significant difference in average MAIA scores between the two groups (W = 333, p = 0.132, CI95% = −0.100–0.750, M_Blind_ = 3.086, M_Sighted_ = 2.736), indicating that the blind group and the sighted control group did not differ significantly in interoceptive sensibility, as measured by questionnaire ratings.

Furthermore, there was no significant difference in the confidence ratings between the blind group and the sighted group (t(44) = 0.411, p = .683, CI95% = −0.991–1.498, M_Blind_ = 5.471, M_Sighted_ = 5.725).

Taken together, these behavioral results align with those presented in Radziun et al. (2023b), which partially relied on the same data. Moreover, they substantiate and confirm the significant differences that we found in cardiac interoceptive accuracy in the current groups of blind and sighted participants who underwent structural MRI scans.

### Structural MRI results

Blind individuals showed increased cortical thickness in the occipital cortex bilaterally, encompassing probabilistic visual areas V1, V2, V3, and V4 (Glasser et al., 2016; whole brain analysis: left cluster size: 4652.66 mm^2^, p = .002; right cluster size: 5747.87 mm^2^, p = .002; Figure 1). This finding is in line with previous studies that have reported thicker occipital cortices in blind compared to sighted individuals (Anurova et al., 2015; Jiang et al., 2009; Park et al., 2009; Voss & Zatorre, 2012).

Similar results were obtained when we included interoceptive accuracy in the analyses as a covariate; that is, we observed increased cortical thickness for blind compared to sighted individuals within probabilistic V1, V2, V3, and V4 (whole-brain analysis: left cluster size: 4522.64 mm^2^; p = .004; right cluster size: 5534.34 mm²; p = .002; see Figure S2 in the Supplementary Material). Thus, as expected, there are differences in cortical thickness between the two groups that cannot be explained solely by differences in interoception.

The region of interest analyses did not reveal group differences outside of the visual cortex. Thus, no significant group differences were observed within cortical areas that have previously been shown to be involved in cardiac interoception (e.g., insula or anterior cingulate cortex; see Schulz, 2016).

When we looked for interactions between group, cortical thickness, and interoceptive accuracy, no clusters survived corrections over the whole brain. However, when we restricted this interaction analysis to our regions of interest, we observed that blind but not sighted individuals showed a significant positive correlation between interoceptive accuracy and cortical thickness in the left and right visual cortices, including probabilistic V1, V2, V3, and V4 (left: r² = 0.40; cluster size: 1573.10 mm², p = .006; right: r² = 0.36, cluster size: 953.68 mm², p = .038; Figure 2). Thus, blind individuals with thicker occipital cortices were more accurate in the heartbeat counting task. No significant interactions between group, cortical thickness or interoceptive accuracy were observed in regions that have been shown to be involved in cardiac interoception.

**Figure 2.**
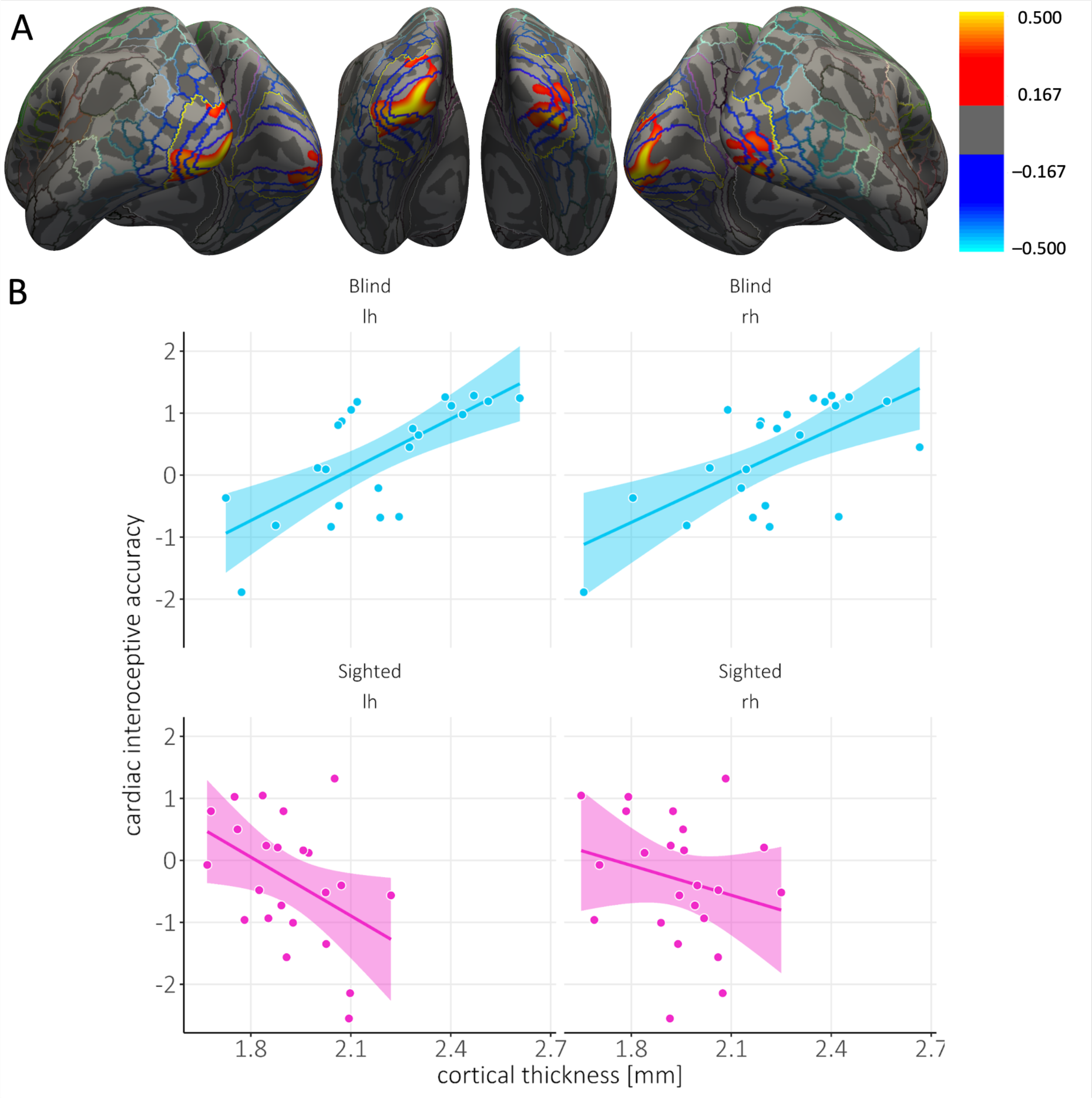
Congenitally blind individuals but not sighted ones showed a positive correlation between visual cortical thickness and interoceptive accuracy. **A** Thresholded maps of correlation coefficients for cortical thickness with cardiac interoceptive accuracy in the group of congenitally blind individuals (CB group, n = 23), in clusters where we observed significant interactions between group, cortical thickness and interoceptive accuracy. Maps are superimposed on the inflated surface (dark gray: sulci, light gray: gyri) of the FreeSurfer standard brain. Colored lines indicate parcellations of the HCP-MMP1.0 atlas (Glasser et al. 2016). The yellow demarcates the regions of interest for the visual cortex and brain areas that have been shown to be associated with cardiac interoception. Clusters that show a positive correlation between cortical thickness and interoceptive accuracy are shown in red/yellow. **B** Average cortical thickness extracted from the significant cluster plotted against the accuracy score from the cardiac interoception task as a function of group (blind: cyan, sighted: pink) and hemisphere. Each dot represents one participant. The line represents the line of best fit. lh = left hemisphere, rh = right hemisphere.

Lastly, we looked for correlations between cortical thickness and interoceptive accuracy in the two groups separately. When we corrected over the whole brain, blind individuals showed a positive correlation between cortical thickness and interoceptive accuracy within the right visual cortex, including V1, V2, V3, and V4 (r² = 0.49, cluster size = 1347.95 mm², p = .035; see Figure 3). In addition, the region of interest analyses revealed significant correlations between cortical thickness and interoceptive accuracy within both the left (r² = 0.55, cluster size = 1248.83 mm², p = .012) and the right visual cortices (r² = 0.50, cluster size = 1256.65 mm², p = .018; see Figure S3 in Supplementary Material). No significant associations between cortical thickness and interoceptive accuracy were observed in regions that have been shown to be involved in cardiac interoception, such as the insula and anterior cingulate cortex.

**Figure 3.**
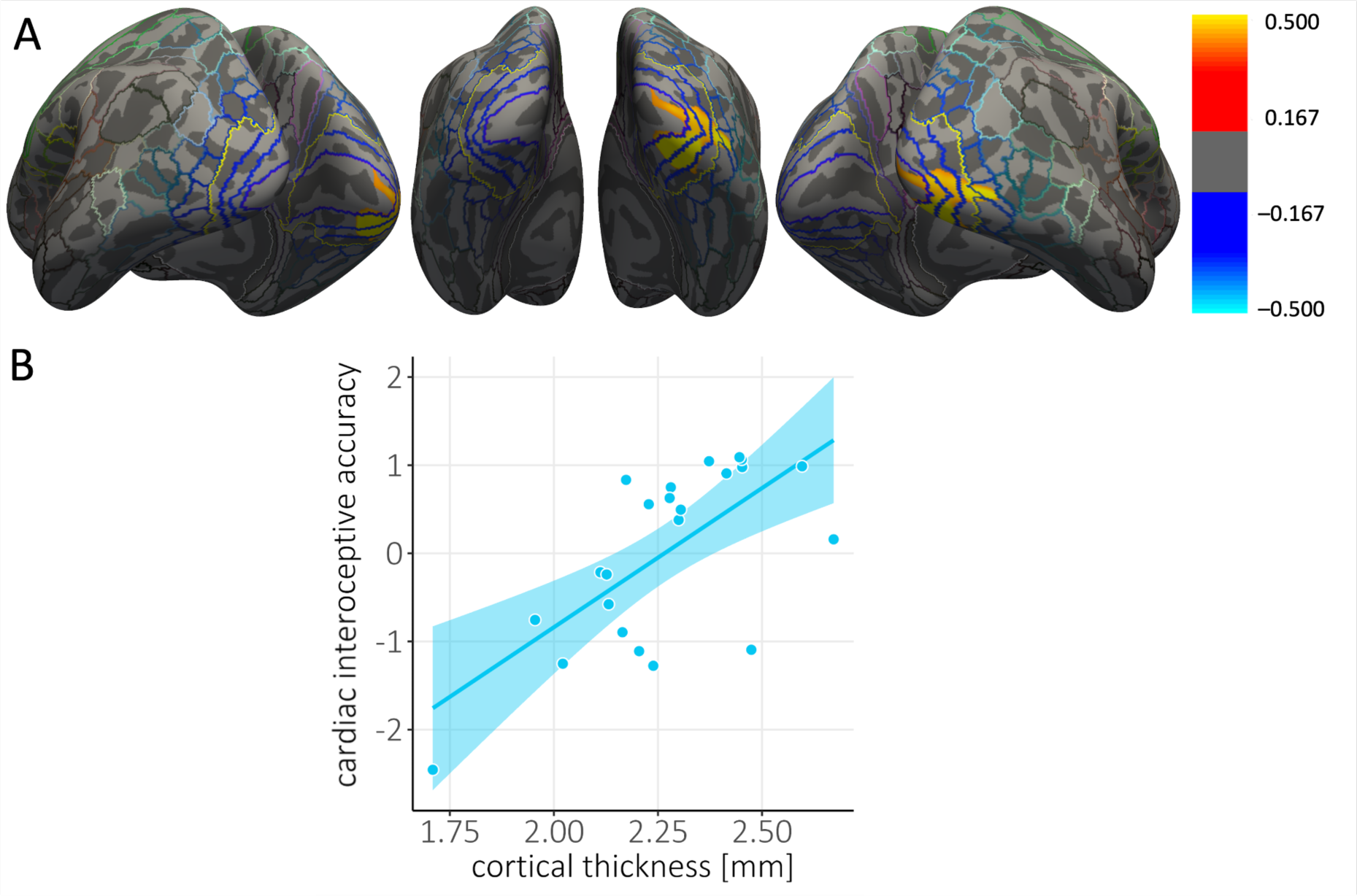
Positive correlation between visual cortical thickness and interoceptive accuracy in congenitally blind individuals. **A** Thresholded maps of correlation coefficients for cortical thickness with cardiac interoceptive accuracy in the group of congenitally blind individuals (CB group, n = 23). Maps are superimposed on the inflated surface (dark gray: sulci, light gray: gyri) of the FreeSurfer standard brain. Colored lines indicate parcellations of the HCP-MMP1.0 atlas (Glasser et al. 2016). The yellow outline demarcates the visual cortex and regions of the brain that have been shown to be associated with cardiac interoception. Clusters that show a positive correlation between cortical thickness and interoceptive accuracy are shown in red/yellow. **B** Average cortical thickness extracted from the significant cluster plotted against the accuracy score from the cardiac interoception task. Each dot represents one participant. The line represents the line of best fit.

**Figure 4.**
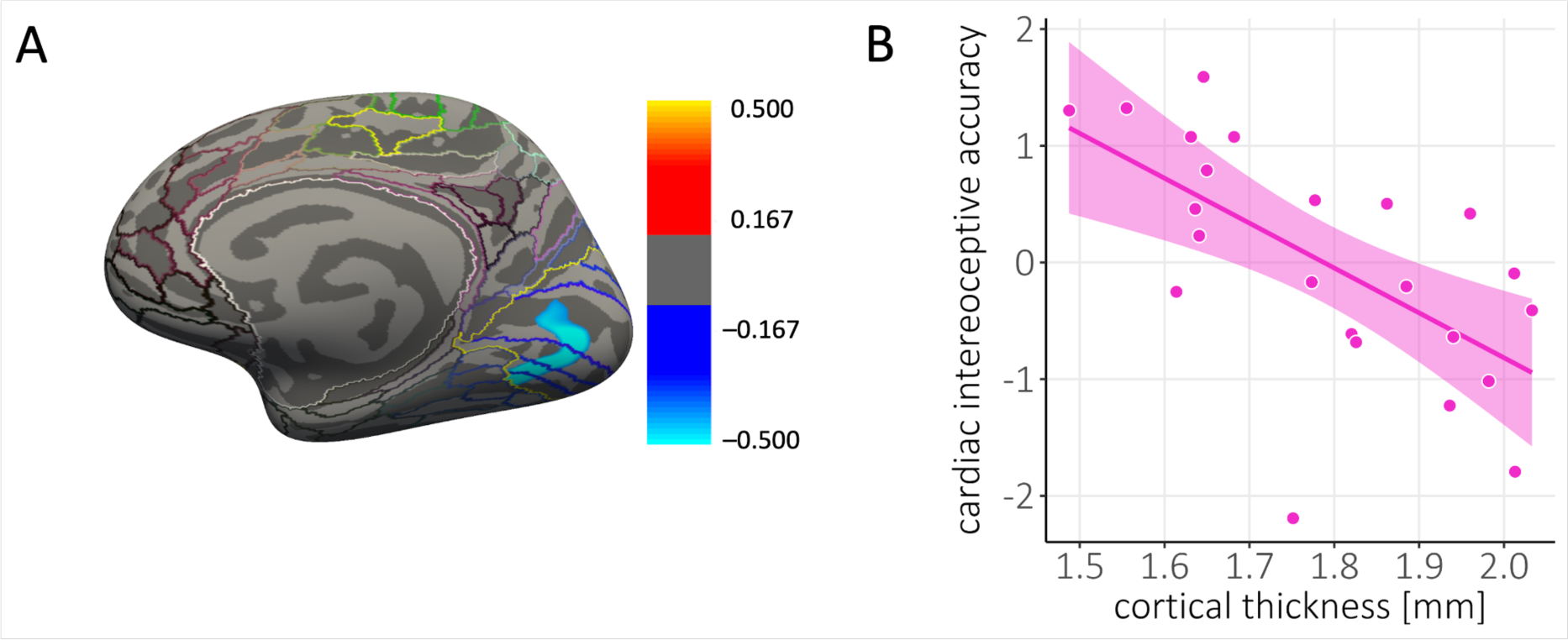
Negative correlation between visual cortical thickness and interoceptive accuracy in sighted individuals. **A** Thresholded maps of correlation coefficients for cortical thickness with cardiac interoceptive accuracy in the group of sighted individuals (SC group, n = 23). Maps are superimposed on the inflated surface (dark gray: sulci, light gray: gyri) of the FreeSurfer standard brain. Colored lines indicate parcellations of the HCP-MMP1.0 atlas (Glasser et al. 2016). The yellow outline demarcates the visual cortex and regions of the brain that have been shown to be associated with cardiac interoception. Clusters that show a positive correlation between cortical thickness and interoceptive accuracy are shown in red/yellow. **B** Average cortical thickness extracted from the significant cluster plotted against the accuracy score from the cardiac interoception task. Each dot represents one participant. The line represents the line of best fit.

When we looked for correlations between cortical thickness and interoceptive accuracy in sighted individuals in a whole-brain analysis, we did not find any significant clusters. Restricting the analysis to our regions of interest, however, revealed a significant cluster in the right visual cortex (r² = −0.50, cluster size = 890.53 mm², p = .036; see Figure 7). Here, sighted individuals showed a negative correlation between cortical thickness and interoceptive accuracy. No associations between cortical thickness and interoceptive accuracy were observed in regions that have been shown to be involved in cardiac interoception.

## Discussion

We investigated whether occipital cortical thickness is related to heightened cardiac interoceptive abilities in congenitally blind individuals. Our results showed that congenitally blind people with thicker occipital cortices have enhanced cardiac interoceptive accuracy. The opposite pattern was observed in sighted individuals; that is, sighted individuals with increased occipital cortical thickness had lower cardiac interoceptive accuracy. Previously, it has been suggested that thicker occipital cortices in blind individuals may reflect atrophy in deafferented structures. However, our finding of a systematic positive relationship between performance on the heartbeat counting task and occipital cortical thickness in blind individuals challenges this notion. Our interpretation is that the behavioral enhancements in heartbeat sensing ability are mediated through cross-modal compensatory plasticity, whereby a thicker occipital cortex provides a behavioral advantage in processing information related to cardiac interoception. This finding is conceptually important because it suggests that cross-modal plasticity following blindness extends beyond exteroception to interoception, encompassing the sense of one’s inner self.

The finding of thicker occipital cortex in the group of congenitally blind individuals compared to the group of sighted controls is in line with previous reports (Aguirre et al., 2016; Anurova et al., 2015; Bridge et al., 2009; Hasson et al., 2016; Jiang et al., 2009; Park et al., 2009; Voss & Zatorre, 2012). However, while numerous studies have reported an increase in cortical thickness in blind individuals, the mechanisms causing this increase are still a matter of debate. MRI studies have reported that after an initial increase in thickness, the cortex appears to thin during development (see Gilmore et al., 2018 for a review; Walhovd et al., 2016; Wang et al., 2019). Similarly, synaptic density in the occipital cortex usually reaches its maximum in the first year after birth and then gradually decreases until it reaches the level seen in adults (Huttenlocher et al., 1982). This process is referred to as synaptic pruning (see Sakai, 2020). It has been hypothesized that cortical thinning is related to the elimination of synapses, since both processes seem to follow a similar time course. Whereas the initial increase in synaptic density does not seem to depend on visual experience, the subsequent pruning of synapses does (Bourgeois & Rakic, 1996). This notion is further supported by studies showing that the cortical thickness of the occipital cortex is related to the age of blindness onset (Li et al., 2017). Based on these findings, it has been hypothesized that increased cortical thickness of the occipital cortex in blind individuals is the result of a disruption of synaptic pruning (Jiang et al., 2009).

However, recent evidence from quantitative MRI and diffusion MRI challenges the notion of cortical thinning during development and instead suggests that the adjacent white matter of the cortex becomes more myelinated (Natu et al., 2019; Whitaker et al., 2016). Thus, it could be hypothesized that what previous studies have identified as thicker occipital cortices in blind individuals reflects persisting immature features and, thus, less myelination of these regions in blind adults. These mechanisms are not mutually exclusive, and a combination of disrupted synaptic pruning and reduced myelination may result in (apparent) thicker cortices in blind individuals.

We observed that the cortical thickness of the occipital cortex was positively correlated with performance on the cardiac interoception task in the group of congenitally blind individuals, but not in the group of sighted controls. These results provide further evidence for the notion that cross-modal plasticity in blind individuals may underlie behavioral enhancements observed in this population (Voss & Zatorre, 2012). The occipital cortex has been shown to be activated to a greater extent in congenitally blind individuals compared to sighted individuals during various non-visual tasks (Amedi et al., 2003; Kujala et al., 1995; Röder et al., 2002), and activation of the occipital cortex and behavioral performance on verbal and auditory tasks has been shown to be correlated in blind individuals (Gougoux et al., 2005, Amedi et al., 2003). Moreover, disruption of the occipital cortex by means of TMS has been shown to impair Braille reading (Cohen et al., 1997; Kupers et al., 2007), verbal processing (Amedi et al, 2004) and sound localization (Collignon et al., 2009), lending further evidence to the notion that the involvement of the occipital cortex in these tasks is functionally relevant. So far, no fMRI studies have investigated the neural correlates of cardiac interoception in congenitally blind individuals. Future fMRI experiments are needed to test the hypothesis that the occipital cortex is involved when blind individuals perform the heartbeat counting task. It will also be important to determine whether different parts of the visual cortex are involved in auditory, tactile, and interoceptive processing, or if the same active regions of the occipital cortex are engaged in different tasks and different sensory processes.

In sighted individuals, performance on tasks involving cardiac interoception has been consistently linked to activation of regions related to the processing of visceral interoceptive signals, such as the insular cortex (Cameron & Minoshima, 2002; Critchley et al., 2004; Herman et al., 2021; Pollatos et al., 2007; Stern et al., 2017; Zaki et al., 2012) and the anterior cingulate cortex (Caseras et al., 2013; García-Cordero et al., 2016; Khalsa et al., 2009; Kleckner et al., 2017). However, in our study, no associations were observed between cardiac interoception and cortical thickness in these regions, for both blind and sighted participants. Although negative findings in neuroimaging studies should be interpreted with caution, this observation speaks against intra-modal plasticity being a critical factor in explaining the superior performance of blind individuals in heartbeat counting tasks. Instead, these negative results together with our positive finding regarding the link between cardiac interoceptive accuracy and occipital cortical thickness points towards cross-modal plasticity being the driving force behind enhanced heartbeat awareness in blind individuals.

Alternatively, it could be hypothesized that the occipital cortex supports cognitive processes associated with good performance in heartbeat counting tasks. For example, it has been suggested that participants may perform this task by estimating rather than counting their felt heartbeats (Desmedt et al., 2020). However, the supporting evidence for this is inconclusive (Schulz et al., 2021; Desmedt et al., 2023; Schulz & Vögele, 2023; Desmedt et al., 2023). Importantly, there is no evidence of differences between blind and sighted individuals in time estimation abilities (Bottini et al., 2015). Our correlative findings cannot resolve the question of whether other cognitive factors may play a role in the observed result. Future fMRI studies could investigate this by examining functional connectivity between the occipital cortex and brain regions that process afferent signals from the heart, exploring possible heartbeat-evoked neural responses in the occipital cortex in blind individuals, and examining how these may relate to enhanced cardiac interoceptive accuracy.

Intriguingly, our results have revealed a negative correlation between occipital cortical thickness and cardiac interoceptive accuracy in sighted individuals. In their seminal paper, Critchley and colleagues (2004) show that attention to cardiac signals decreases activity in occipital areas in sighted participants. This is in line with the cross-modal deactivations of the occipital cortex that have been reported when participants attend to and perform tasks in other sensory modalities (Kawashima et al., 1995; Limanowski et al., 2020; Morita et al., 2019). By contrast, Herman, Palmer, Azevedo, & Tsakiris (2021) have demonstrated that attention to and detection of interoceptive signals can lead to increased activations of the occipital cortex, potentially pointing to a functional role of the occipital cortex during interoceptive tasks. The exact relationship between visceral signal processing and the occipital cortex in sighted individuals remains to be clarified (see Azzalini, Rebollo, & Tallon-Baudry, 2019). However, if the occipital cortex indeed plays a role in processing interoceptive signals in sighted individuals, it could be hypothesized that our results reflect potentiation of pre-existing architecture that has the necessary representational and computational capacity for processing cardiac interoceptive signals (Makin & Krakauer, 2023; Meredith et al., 2011).

The limitations inherent in relying on gray matter thickness as a proxy for microstructural changes should be acknowledged. While cortical thickness serves as a valuable metric, it provides a macroscopic view that lacks the specificity required to elucidate the intricate neurophysiological adaptations occurring within deprived occipital regions. This study, therefore, prompts consideration of alternative methodologies, such as quantitative MRI or postmortem anatomical work in blind individuals in order to complement and extend our findings. These approaches may offer a more nuanced understanding of the synaptic and myelin-related adaptations that contribute to the observed changes in cortical thickness, ultimately unraveling the complex interplay between anatomical alterations and heightened perceptual abilities.

In conclusion, we have conducted the first study investigating the link between changes in cortical thickness and blind individuals’ ability to sense their own heartbeats. Our results suggest that structural plasticity in the occipital cortex of congenitally blind individuals supports the enhanced processing of cardiac interoceptive signals in a heartbeat counting task. This observation advances our understanding of the link between structural changes and behavioral enhancements after blindness and suggests that such cross-modal plasticity extends to the processing of signals from the body’s inner organs, thus expanding our understanding of the limits of cross-modal plasticity in blindness.

## Acknowledgements

We would like to thank Cordula Hölig for her advice on the data analysis pipeline, as well as all the participants, without whom this research would not have been possible.

## Authorship contribution statement

Anna-Lena Stroh and Dominika Radziun served as lead for conceptualization, investigation, methodology, writing–original draft, and writing–review and editing. Maksymilian Korczyk served in a supporting role for data curation, project administration, and writing–review & editing. Laura Crucianelli served in a supporting role for methodology and writing–review and editing. Marcin Szwed and H. Henrik Ehrsson served as lead for supervision and served in a supporting role for conceptualization and writing–review and editing. Marcin Szwed and H. Henrik Ehrsson contributed to funding acquisition and resources equally.

