## Supplementary material for "Blind individuals’ enhanced ability to sense their own heartbeat is related to the thickness of their occipital cortex"

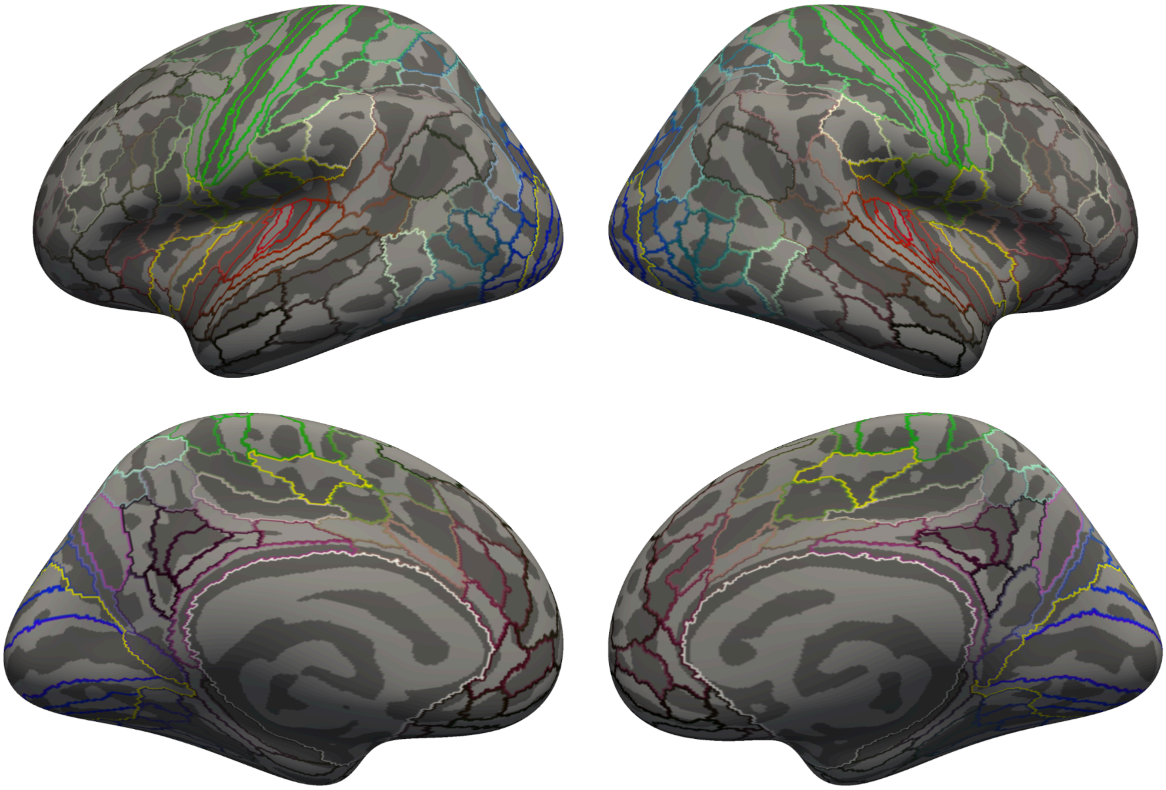

**Figure S1.** Inflated surface (dark gray: sulci, light gray: gyri) of the FreeSurfer standard brain. Colored lines indicate parcellations of the HCP-MMP1.0 atlas (Glasser et al. 2016); the yellow line demarcates the visual cortex and regions that have been shown to be involved in cardiac interoception (Schulz et al., 2016).

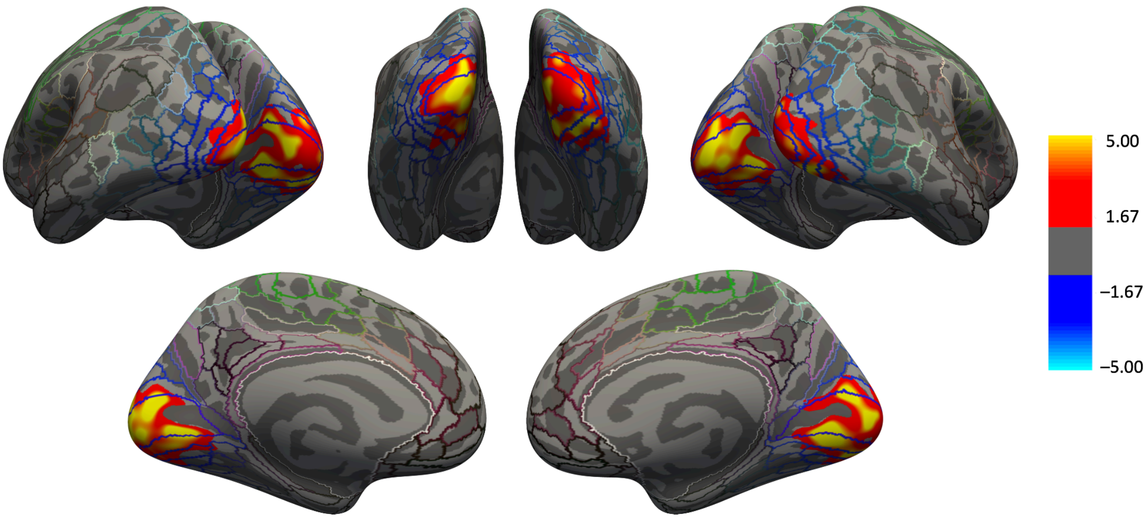


**Figure S2.** Group differences in visual cortical thickness with interoceptive accuracy as a covariate. **A** Thresholded statistical significance maps (vertex-wise p < .01, cluster-wise p < .05, two-sided) display cortical thickness differences between congenitally blind individuals (CB group, n = 23) and sighted controls (SC group, n = 23). Maps are superimposed on the inflated surface (dark gray: sulci, light gray: gyri) of the FreeSurfer standard brain. Colored lines indicate parcellations of the HCP-MMP1.0 atlas (Glasser et al. 2016). Clusters with higher cortical thickness in the CB group are marked in red.
